# Real time health monitoring through urine metabolomics

**DOI:** 10.1101/681742

**Authors:** Ian Miller, Sean R. Peters, Katherine A. Overmyer, Brett R. Paulson, Michael S. Westphall, Joshua J. Coon

## Abstract

Current healthcare practices are reactive and based on limited physiological information collected months or years apart. By enabling patients and healthy consumers access to continuous measurements of health, wearable devices and digital medicine stand to realize highly personalized and preventative care. However, most current digital technologies provide information on a limited set of physiological traits, such as heart rate and step count, which alone offer little insight into the etiology of most diseases. Here we propose to integrate data from biohealth smartphone applications with continuous metabolic phenotypes derived from urine metabolites. This combination of molecular phenotypes with quantitative measurements of lifestyle reflect the biological consequences of human behavior in real time. We present data from an observational study involving two healthy subjects and discuss the challenges, opportunities, and implications of integrating this new layer of physiological information into digital medicine. Though our dataset is limited to two subjects, our analysis (also available through an interactive <u>web-based visualization tool</u>) provides an initial framework to monitor lifestyle factors, such as nutrition, drug metabolism, exercise, and sleep using urine metabolites.

## Introduction

Current medical practice is reactive. Annual checkups measure only a few basic phenotypes and often fail to predict serious health threats such as cancer, dementia, or exposure to pathogens. Instead, most disease is not detected until critical symptoms present, which is often too late for meaningful or cost-effective intervention. Owing to this lack of data, the current model of healthcare is periodic and geared to manage disease symptoms at their onset rather than preventing or reversing the underlying etiology. Humans would undoubtedly benefit from integrated technology to quantify and monitor deviations from baseline wellness using physiological phenotypes^1^. Yet access to actionable information on personal physiological health remains limited.

There are currently two avenues for continuous monitoring of health and disease: (1) consumer-grade wearables and (2) clinical-based precision medicine. Wearable devices such as smart watches are broadly accessible and increasingly popular as consumer products. Data from these devices has the advantage of being continuously and passively collected from large populations of people. Many companies have since devoted significant resources to leverage tools in big data and artificial intelligence (AI) to provide actionable insights from these popular products. For instance, Apple (CA, USA) has recently received FDA approval to provide users with alerts to detect atrial fibrillation^2^. This diagnostic capability was made possible by widespread consumer participation, which provided expansive datasets to train AI models. The Apple Heart Study involved roughly 400,000 participants and models constructed from this initial dataset were validated with a clinical trial involving approximately 600 participants.^2^

Given sufficiently large datasets, heart rate information alone can suggest the onset of diverse disease processes^3^. However, this type of data offers little information on the origins, mechanisms, and progression of disease. For instance, while an elevated resting heart rate may indicate a number of adverse medical events, including an infection^3^, such data is not able to distinguish between bacterial and viral infections. This lack of mechanistic information leaves patients and health care providers unable to implement targeted therapeutic intervention and, in this case, antibiotic stewardship.

On the other end of the spectrum of longitudinal monitoring are tools for clinically-based precision medicine. These include deep genome sequencing and integration with multidimensional clinical phenotypes such as transcriptomics, proteomics, metabolomics, and metagenomics datasets. There are a number of large-scale efforts underway to provide multi-omic phenotyping for large cohorts, such as the Pioneer 100 Wellness Project^1,4^ and the NIH All of Us program^5^. While these initiatives have proven to successfully leverage diverse physiological datasets to enable meaningful intervention^1^, they remain hampered by their relative inaccessibility and periodic nature. In other words, while high quality data provides clinically actionable insights, it is expensive, invasive, and difficult to collect, resulting in collections on the scale of months rather than days^1^.

As Leroy Hood and others have long proposed, modern medicine will only be truly effective once it is has transitioned from reactive disease care to a framework that is “predictive, preventive, personalized, and participatory”^6^. To combine the accessibility of wearable devices with the robustness and quality of clinical medicine, a third option is needed; one that provides quantitative measurements of health and mechanistic insights into the origins and progression of disease. We hypothesize that realtime metabolic phenotyping (i.e., “metabolomics”) using urine could fill this void by providing a quantitative fingerprint of metabolic health along with information about exposure to toxins, drugs, and pathogens^7^. In theory, continuous metabolic measurements could be collected at home and in the workplace, providing molecular insights into underlying disease processes, such as distinguishing between patients with related strains of infectious bacteria^8^, as well as quantifying the effect of lifestyle decisions on health and disease. Lifestyle factors such as nutrition, alcohol and tobacco usage, sleep, and physical activity are well known to contribute to the risk of developing chronic disease, which costs the United States alone $2.97 Trillion a year, or 90% of all healthcare expenditures.^9^ By empowering consumer participation with actionable information and the classification of disease using a continuum of molecular phenotypes rather than discrete clinical symptoms^6^, the cost and efficacy of healthcare could be dramatically improved.

While a number of biological matrices, including saliva and blood, could be used as a source of metabolic information, urine offers some key advantages as it can be easily collected passively, non-invasively, and longitudinally^10^. Urine is a rich source of cellular metabolites, most stemming from filtration of blood in the kidneys, which excrete about a half cup of blood every minute^11^. Urine has long been recognized as a rich fluid for medical diagnostics and presently many clinical assays are performed on this biological fluid^12–14^. Approximately 4,500 metabolites have been documented in urine^13,15^, showing connections to approximately 600 human conditions^12,13^ including but not limited to: obesity^16^, cancer^17^, inflammation^18^, neurological disease^19^, and infectious disease^8^. Further, pregnancy, ovulation, urinary tract infection, diet, and exercise induce metabolomic signatures that can be observed in urine^14^. Finally, many drugs and their metabolites are readily detected from urine, presenting the opportunity for dosage tailored to the individual and monitoring compliance as well as effective stratification for clinical trials, which can greatly reduce the cost of pharmaceutical development^6,7^.

Over the course of 10 days, we collected every urine sample from two healthy individuals and tracked hundreds of urine metabolites using gas chromatography and mass spectrometry (GC-MS) along with other biometric data provided by nutritional and fitness smartphone applications. Though other studies have measured the concentrations of urine metabolites from larger populations over time with other physiological phenotypes^14,20^, we are unaware of any studies with the time resolution or smartphone data integration we present here. Our aim was to explore this combination of smartphone and metabolomics data as a means to understand the biological consequences of lifestyle in real time.

## Methods

### Sample Collection

Urine samples were collected midstream and volume was measured using sterile 500 mL plastic beakers from which urine was then decanted into cups provided in the BD Vacutainer® Urine Complete Cup Kit. Samples were then transferred into 8 mL urinalysis plus conical urine tubes. These vacuum sealed 8 mL tubes were either immediately deposited into −80 °C freezers or temporarily stored in dry ice using portable coolers overnight before being deposited into −80 °C freezers. Quantitative values for sample volume along with sample collection times are available in **Supplementary Dataset 3**.

### Ethics approval and consent to participate

The University of Wisconsin - Madison Institutional Review Board was consulted prior to collecting data for this study. The IRB determined review was not required for this study because no human subjects were enrolled.

### Sample preparation and analysis

All urine samples were prepared by sampling 100 µL into Thermo Scientific 300 µL Amber Vials with inserts and subsequently evaporated to dryness using a Thermo Scientific SpeedVac® Concentrator. Samples were then derivatized for gas chromatography analysis using 50 µL solution of 1:1 pyridine: N-Methyl-N-(trimethylsilyl)trifluoroacetamide with 1% trimethylchlorosilane (chemicals obtained from Sigma Aldrich) and incubated at 60 °C for 30 minutes. Samples were then injected onto a Thermo Scientific Gas Chromatography-Fourier Transform Mass Spectrometry (GC-FTMS) Orbitrap using a temperature gradient starting at 100 °C (hold time of one minute), and increasing at a rate of 8.5 °C per minute until reaching 260 °C. The temperature gradient rate was then increased to 50 °C per minute until reaching a final temperature of 320 °C (hold time of four minutes). Split ratio was set to 10:1 with a carrier gas flow of 1.200 mL/min. The MS transfer line and ion source temperatures were set to 300 °C and 250 °C, respectively. The instrument scanned in Full MS-SIM mode at 30,000 resolution. The AGC target was set to 1.0e6 with a scan range of 50 to 650 m/z. Ionization mode was set to electron ionization (EI).

Raw files were subsequently processed using in-house software^21^ for deconvolution, peak alignment, quantitation, and identification. Cutoffs for peak quantitation were set to a minimum fragment count of 10, minimum observation of a given peak across all files set to 33%, and analyte/background signal set to 10. Spectra were then matched against the unit resolution library curated by the National Institute of Standards and Technology (NIST), and a high resolution library developed in house in collaboration with Thermo Scientific.

### Preparation and analysis of ethyl glucoronide standard

Ethyl glucuronide standard (100 ug/mL, Sigma Aldrich) was dried down in an Amber autosampler vial using a SpeedVac® Concentrator. This standard was derivatized and analyzed as described above.

### Biometric data collection

Nutritional data was recorded daily using the Lose It! App. For Subject 1, active calories were recorded with an Apple Watch Series 2 (Model A1758, software version 4.3.2 (15U70)) and hours of sleep were calculated using the Sleep Cycle App. Summary statistics are available in **Supplementary Table 1**.

### Analysis of the Human Urine Metabolome Database (HMDB)

The XML version of the human urine metabolome database (urine_metabolites.xml) was downloaded on October 7, 2018. The various aggregated files and scripts used in this analysis are accessible in the associated GitHub repository (see **Code Availability**) and as **Supplementary Datasets 1-2**. For the purposes of visualization, broader disease categories, such as “Cancer” and “Inflammation” were curated manually (**Supplementary Dataset 2**). A treemap (**Fig. 2c**) was generated using the ggplot2^22^ and treemapify^23^ modules in R.

**Fig. 1.**
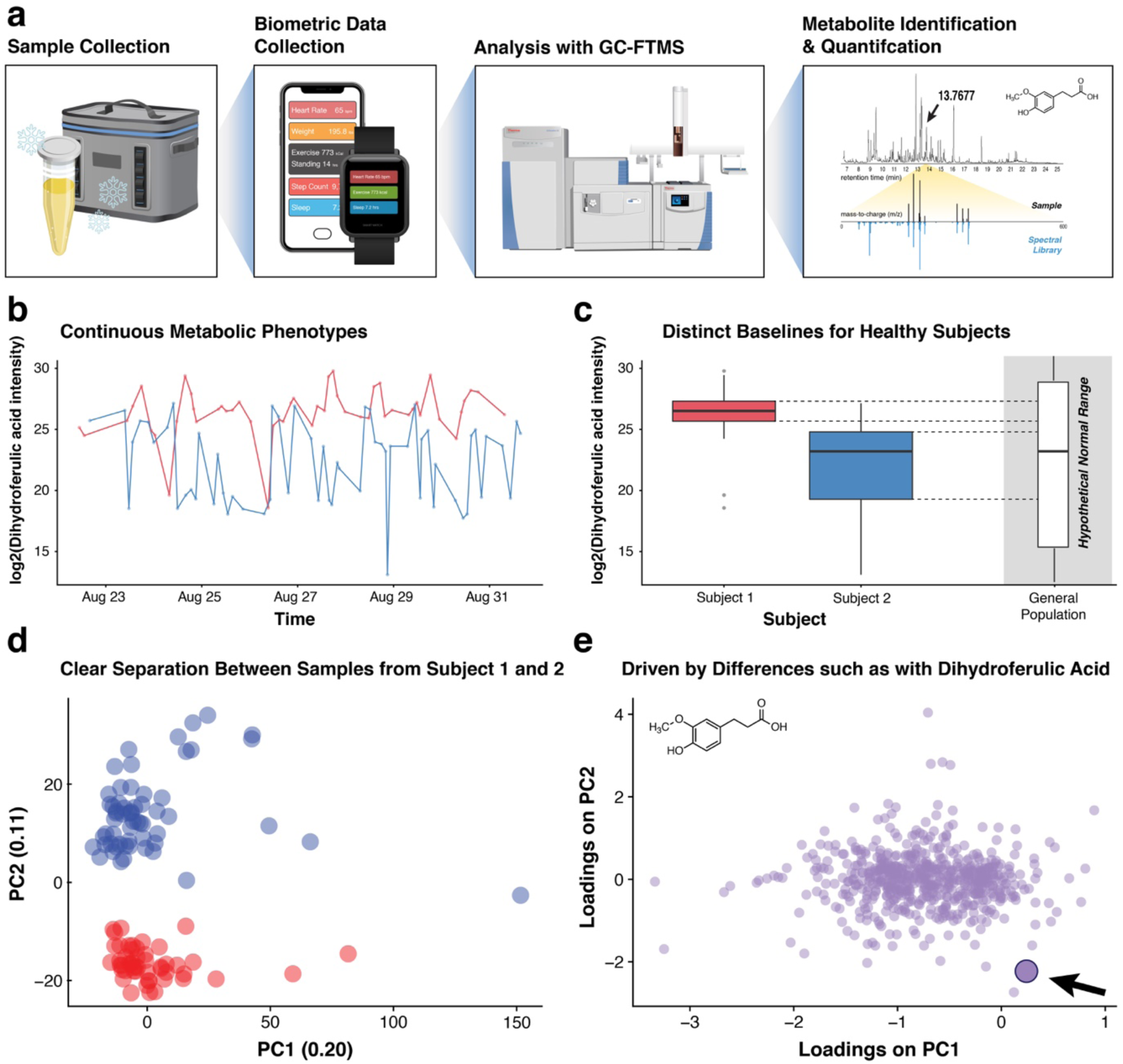
GC-MS metabolomics provides continuous and distinct metabolic phenotypes for two healthy individuals. (**a**) Every urine sample (109 total; Subject 1 [red], *n* = 50; Subject 2 [blue], *n* = 59), along with biohealth data from smartphone applications, was collected for 10 days. Samples were dried down, derivatized with n-methyl-n-(trimethylsilyl)trifluoroacetamide, and analyzed with high resolution GC-Fourier Transform Mass Spectrometry (FTMS). (**b**) Deconvolution and quantification with in-house software^21^ provide time series profiles for 603 metabolite features. (**c**) The log2-intensity of a metabolite feature identified as dihydroferulic acid, a known urine metabolite, revealed different baseline concentrations for Subject 1 and Subject 2, compared to a hypothetical range for the general population. (**d**) Scores plot from principal component analysis (PCA) based on log2-normalized intensity values shows clear separation between Subject 1 and Subject 2. Each point represents a sample and is colored by Subject. (**e**) PCA loadings plot where each point represents a metabolite feature. An interactive version of (**d**) and (**e**) are provided in the companion webtool.

**Fig. 2.**
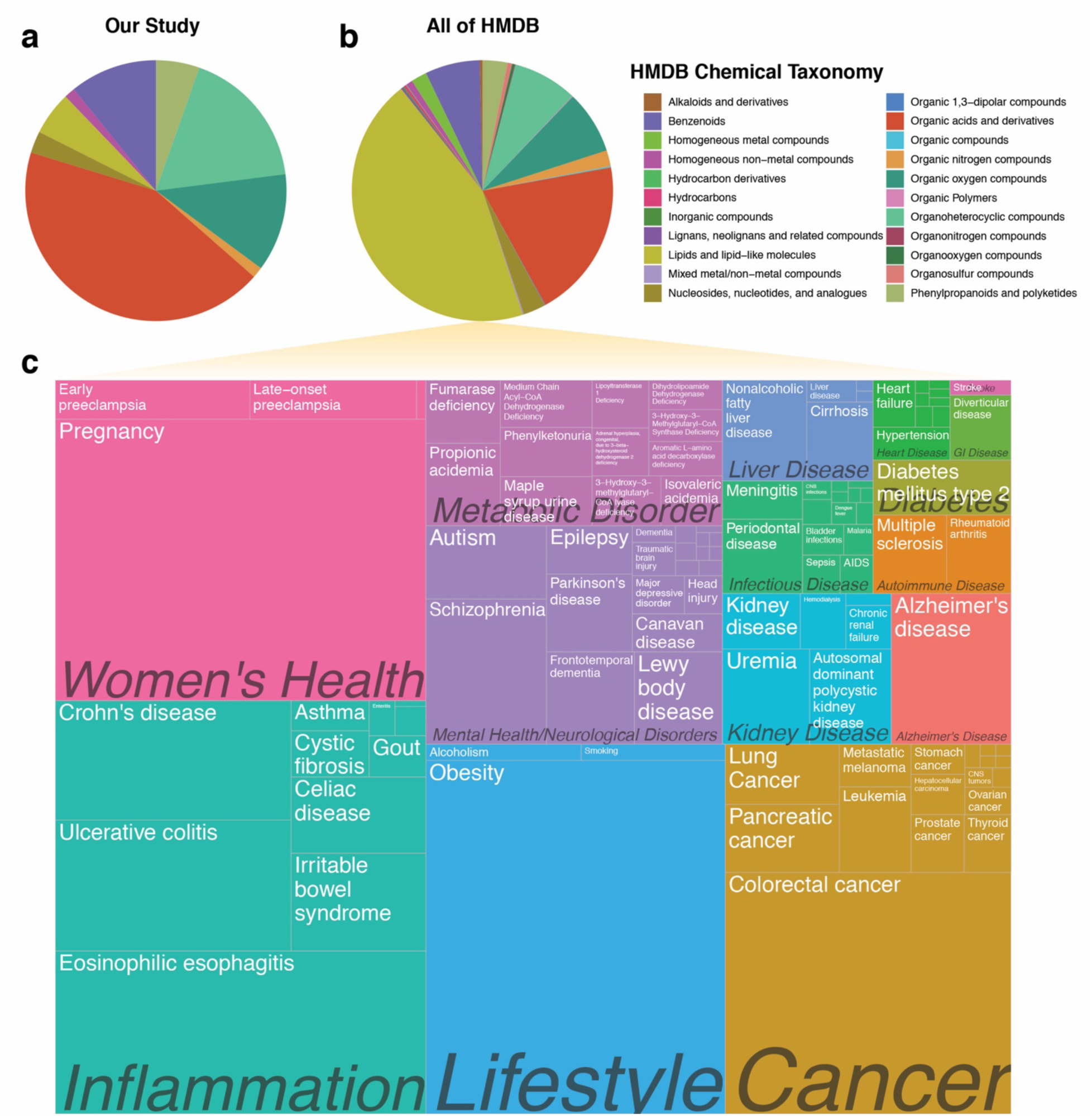
There 4240 known urine metabolites in the human metabolome database (HMDB), 1424 of which have literature associations to a diverse set of human conditions. (**a**) Pie chart based on counts of HMDB chemical taxonomy (“Super Class”) for metabolites in this study. (**b**) Pie chart based on counts of each HMDB chemical taxonomy for all urine metabolites in HMDB. (**c**) Treemap of diseases, scaled by number of metabolites with HMDB literature associations.

### Statistical analysis

Metabolite intensity values were normalized to the total ion current (TIC) for the RAW files for each sample using previously described software^21^. Principal component analysis was conducted on log2-transformed TIC-normalized intensity values using the decomposition.PCA() method in Python’s skickit learn module^24^. Welch’s Two Sample t-test was performed in R using the t.test(var.equal = F) function. Repeated measures correlation^25^ was used to correlate daily log2-transformed average metabolite intensities with biometric values where biometric data were available for both subjects using the rm_corr() function in the Pingouin^26^ package in Python. Skipped (robust) Spearman’s rho was used to correlate log2-transformed daily average metabolite intensities with biometric values for data exclusively available for Subject 1 (i.e., physical activity and sleep) using corr(method=‘skipped’) from Pingouin^26^. *P*-values from the repeated measures correlation, and Spearman’s rho were adjusted for multiple hypothesis testing (where the number of tests was considered the number of metabolite features observed [603]) using the Benjamini Hochberg false discovery rate (FDR) procedure via the fdr(method=‘fdr_bh’) method in Pingouin and are presented as *q*-values in the manuscript. All *p*-values from hypothesis testing are based on two-sided tests and degrees of freedom and/or *n* values are presented as they appear in the **Results**. Further statistical details on correlation results can be found in **Supplementary Dataset 4**.

Linear regression analysis was used to test for diurnal effects on metabolite concentrations. Samples were binned by “morning” (6 am to 12 pm), “afternoon” (12 pm to 6 pm), “evening” (6 pm to 12 am), and “late night” (12 am to 6 am). Only two samples (both from Subject 1) were collected between 12 am and 6 am and were excluded from this analysis under the assumption that they represent outliers and would create an unbalanced group for regression analysis, which was performed using the linear_regression() function in Pingouin^26^. An effect was considered significant if the reported *p*-value associated with the coefficient for the TimeOfDay designation (see **Supplementary Dataset 3**) was less than or equal to an alpha of 0.05.

### Data availability

A companion web tool is available via the following URL:http://3.16.13.214:6004/dash, and provides an interactive visualization of **Fig. 1d, Fig. 1e**, and **Fig. 3b**. RAW data files were uploaded to the MassIVE database https://massive.ucsd.edu/ProteoSAFe/static/massive.jsp) under the accession number: MSV000083880 (doi:10.25345/C5B33S). Processed and normalized metabolite intensity values along with other relevant metadata are provided in **Supplementary Datasets 1-4** in this study.

**Fig. 3.**
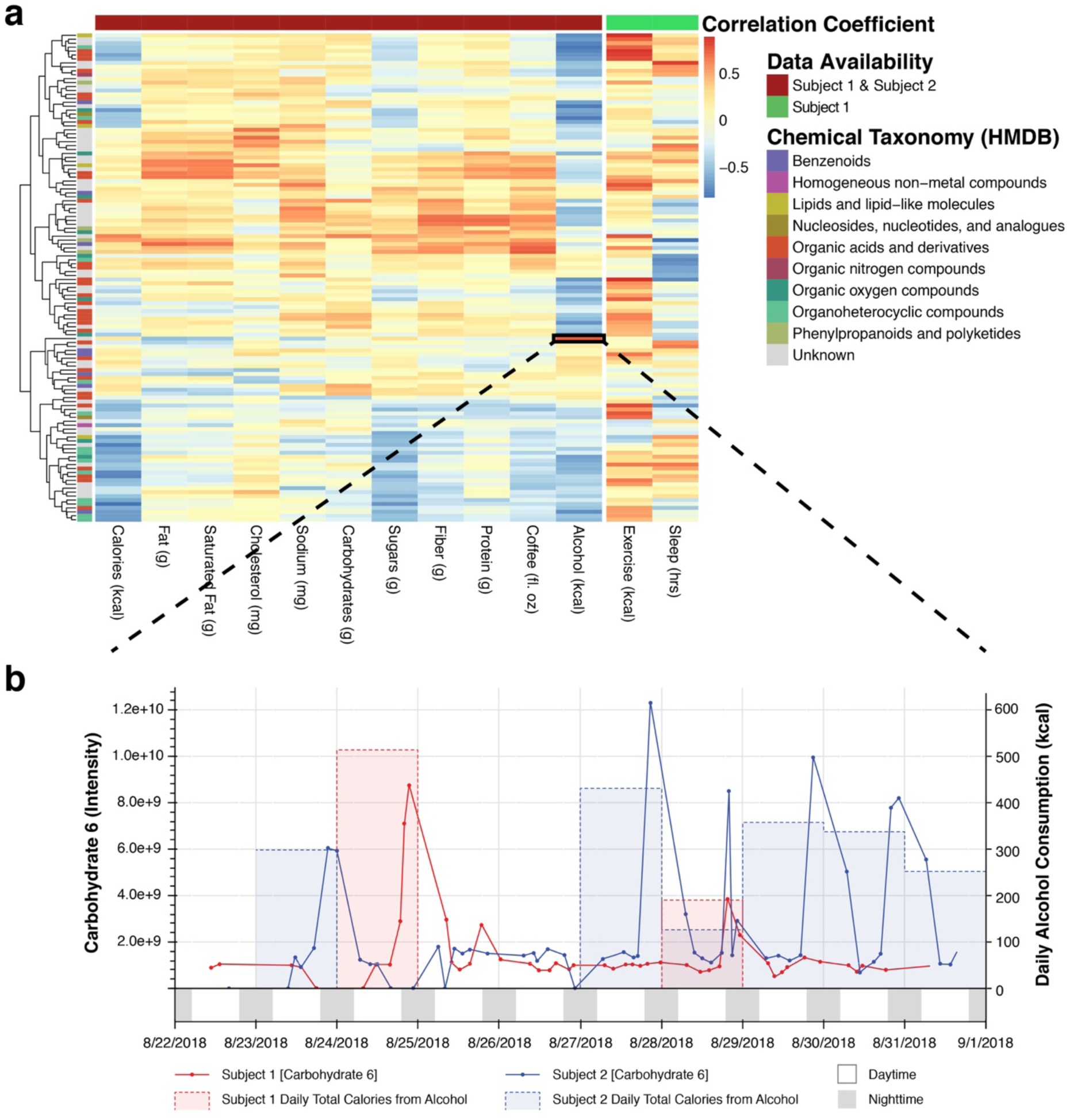
Urine metabolites reflect patterns of health and lifestyle. (**a**) Heatmap whereby each row represents an identified metabolite and each cell represents the strength of correlation between daily average metabolite intensity and a given biometric dataset collected from smartphone apps. Repeated measures correlation^25^ was used where data was available for both subjects whereas Robust Spearman’s Correlation^31^ was used for data only available for Subject 1 (i.e., exercise and sleep). (**b**) An example correlation (r = 0.812, *p* = 1.32e-4, *q* = 0.0121, dof = 14) between alcohol consumption (in kcals) and a carbohydrate compound (“carbohydrate 6”), most likely representing xylitol. An interactive version of (**b**) is provided in the companion webtool.

### Code availability

All source code used for analysis in this study, including for the interactive data visualization tool, is available on github: https://github.com/ijmiller2/RealTimeUrineMetabolomics.

## Results

### Urine metabolites provide distinct and continuous metabolic phenotypes

We collected 109 urine samples (50 for Subject 1 and 59 for Subject 2) over 10 days along with biometric measurements provided by smartphone and smartwatch applications, including those for nutrition, exercise, and sleep (**Fig. 1a** and **Supplementary Table 1**). Urine samples were lyopholized, resuspended, and derivatized with solution containing N-Methyl-N-(trimethylsilyl)trifluoroacetamide, and subsequently analyzed with high resolution Gas Chromatography-Fourier Transform Mass Spectrometry (GC-FTMS). The resulting data were deconvoluted with previously described software^21^, which detected and quantified 603 metabolite features across the 109 individual urine samples. Of these 603 metabolite features, 101 metabolites were annotated/identified based on spectral matching (see **Methods**). An additional 24 features were able to identified as carbohydrates, but were not assigned a specific molecular structure due to the highly similar fragmentation patterns for certain sugars, consistent with a Level 3 assignment according to the standards proposed by the Metabolomics Standards Initiative^15,27,28^.

Each one of these 603 metabolite features provided a continuous metabolic phenotype for both subjects. Compounds, such as dihydroferulic acid–a metabolite of phenolic compounds previously detected in urine^29^–showed different baseline levels for Subject 1 and Subject 2 (**Fig. 1b** and **Fig. 1c**). Though this study was not designed for absolute quantification, the ability to detect different relative baseline metabolite levels for two health subjects (as well as deviations therefrom) offers a proof of principle for continuous urine analysis enabling personalized medicine. For instance, given the higher and tighter distribution of dihydroferulic acid concentrations for Subject 1 compared to Subject 2, deviations from either of these individual baselines would likely be more clinically meaningful than deviations from a hypothetical normal range established for the general population (**Fig. 1c**).

Principal component analysis of the log2-normalized metabolite intensity values showed clear separation between subject samples across PC2 (**Fig. 1d**). An analysis of the corresponding loadings plot of PC2 (**Fig. 1e**) shows dihydroferulic acid, histidine, and phenoxyacetic acid among the top identified metabolite features by rank. While the average volume of samples from Subject 1 was significantly higher than that of Subject 2 (Welch’s Two Sample t-test, t = 7.8739, dof = 81.614, *p* = 1.275e-11), neither PC1 nor PC2 reflect systematic differences in sample volume, run order, or batch number (**Supplementary Fig. 1**), suggesting that the separation in these two planes is driven by biological rather than technical variance. The clear distinction between samples derived from two healthy subjects are not necessarily surprising given that other urine metabolomics studies have shown the ability of unsupervised techniques to resolve a wide range of biological traits and clinical phenotypes^12^.

### Analysis of Literature Disease Associations

Each identified metabolite was searched against the synonyms in the Human Urine Metabolome Database (HMDB) (http://www.urinemetabolome.ca/)^13^. For each metabolite with a matching synonym (75/101), features of interest such as chemical taxonomy and associated diseases (based on previous literature mining efforts^13^) were tabulated (**Supplementary Dataset 1**). These compounds covered a range of chemical classes (**Fig. 2a**), as defined in HMDB as chemical “Super Class,” but had a higher fraction of organic acids compared to metabolite classes represented by the entire HMDB database (**Fig. 2b**). This difference in chemical compositions is likely a result of minimal sample processing and the use of GC-MS^12,13^, which favors volatile and typically lower molecular weight compounds (see **Methods** for more details). In total, 65 metabolites with corresponding HMDB data entries had some type of literature association with diseases, 48 of which had connections to various forms of cancer and 19 with Alzheimer’s Disease (**Supplementary Dataset 1**).

A broader analysis of the 4240 metabolites available in the downloadable database shows that 1424 of these compounds have disease associations in a least one piece of literature. Diseases and conditions ranged from having one (e.g., “Cervical Cancer”) to 586 associated metabolites (“Obesity”) (**Fig. 2c** and **Supplementary Dataset 2**). Although these connections are not clinically validated biomarkers of disease, they may suggest potential for applications of continuous urine analysis in digital health and personalized medicine. In fact, many of the diseases that the Center for Disease Control (CDC) lists as leading causes of death in the United States, such as cancer, diabetes, and kidney disease^30^, have associations to metabolites that can be detected in urine. Collectively the conditions associated with chronic disease create trillions of dollars in cost to the health care system in the United States alone. Many of these diseases are associated with lifestyle factors, such as tobacco, and alcohol usage, and lack of physical activity^30^.

### Metabolite levels reflect nutrition, lifestyle, physical activity, and sleep

To explore connections between urine metabolite concentrations and others measures of health and lifestyle, various biometric data were collected contemporaneously using smartphone applications (see **Methods**). Both subjects collected nutritional data and Subject 1 collected further data on exercise and sleep (**Supplementary Table 1** and **Supplementary Dataset 3**). Due to the disparate time scale for which this data was collected (urine samples were collected four to eight times per day as they were generated, while data such as sleep only provides one time point per day), and to control for diurnal variability (explored further below), daily average metabolite intensities were correlated with biometric values using repeated measures correlation^25^ (where data is available for both subjects) and Robust Spearman’s Correlation^31^ (where data is available only for Subject 1) (see **Methods, Supplementary Dataset 3-4**). Interestingly, there are two predominant groups of metabolites that are (1) positively (see upper half of heatmap) and (2) negatively (see lower half of heatmap) correlated with caloric and nutrient intake (**Fig. 3a**). Perhaps the former represent compounds that are either food-derived or linked to energy metabolism, whereas the latter represent endogenous compounds that belong to metabolic pathways separate from energy metabolism. While an interactive web-based tool to further explore and visualize these correlations is available, (see **Methods** for info on data and source code availability), here, we highlight a few putative connections between metabolite concentrations and over the counter (OTC) medication usage, coffee and alcohol consumption, exercise, and sleep (**Fig. 3**).

Subject 1 drank coffee twice a day (at approximately 7 am and 9 am), and Subject 2 drank coffee every morning around 8 am except for on August 25 and August 27, 2018. These consumption patterns are consistent with our measurements of compounds with known associations to coffee consumption (**Supplementary Fig. 2**). Furoylglycine, a biomarker of coffee consumption^32,33^, and its corresponding intensity was consistent with notes of when coffee was consumed each day (r = 0.617, *p* = 0.011, *q* = 0.201, dof = 14; **Supplementary Fig. 2**). Quinic acid, another known metabolite from coffee^33^, also tracked well with coffee consumption (**Supplementary Fig. 3;** r = 0.787, *p* = 2.93e-4, *q* = 0.0884, dof = 14).

A metabolite that was putatively identified by spectral database searching software (see **Methods** for details on metabolite identification) as a carbohydrate compound (termed “carbohydrate 6” in **Supplementary Datasets 1**,**3**,**4**) was well correlated with alcohol consumption (as measured in kcal) (r = 0.812, *p* = 1.32e-4, *q* = 0.0121, dof = 14, **Fig 3b**). These calories from alcohol include a variety of alcohol types (beer, wine, whiskey, gin, tequila, cognac, and vermouth) and thus are more likely to reflect ethanol consumption rather than a compound specific to a certain type of beverage. Further manual inspection of the spectral matches to this unannotated metabolite feature suggested that it is most likely a sugar alcohol, with the highest dot product score to xylitol. Follow up analysis using a standard of ethyl-glucuronide, an established metabolite and biomarker of ethanol consumption^34^, was then analyzed and added to our in-house spectral database (see **Methods**). A separate (initially unidentified) metabolite feature at a retention time (RT) of 14.842677 and m/z 217.1075485 was subsequently identified as ethyl-gluconoride **Supplementary Fig. 4**). While the repeated measures correlation coefficient for ethyl-glucuronide and alcohol consumption was lower (r = 0.657, *p* = 5.70e-3, *q* = 0.0508, dof = 14) than for the putative sugar alcohol (“carbohydrate 6”), this discrepancy is likely a result of the nature of the metabolite’s pharmacokinetics; ethyl glucuronide has a longer half-life than ethanol^34^.

Subject 2 reported taking acetaminophen at 8:45 pm on August 25, 2018. A spike in an ion intensity consistent with acetaminophen shows a corresponding increase observed in next sample in sequence, which was collected at 7:15 am on August 26, 2018 (**Supplementary Fig. 5**). No such spike was observed for Subject 1, who did not record taking any acetaminophen throughout the collection of these samples.

In addition to nutritional information collected for Subject 1 and Subject 2, data on physical activity and sleep was collected for Subject 1. Hypoxanthine, a degradative purine product resulting from ATP breakdown in muscle tissue during exercise^35,36^, correlated with physical activity (r = 0.833, *p* = 0.0102, *q* = 0.472, *n* = 8; **Supplementary Fig. 6** and **Supplementary Dataset 4**). Sleep was anticorrelated with hydrocaffeic acid (r = −0.857, *p* = 0.0137, q = 0.551, *n* = 8; **Supplementary Fig. 7** and **Supplementary Dataset 4**), a metabolite of caffeic acid^33,37^, which in turn has been shown to affect sleep latency in rats^37^. However, neither of these metabolite correlations passed the significance threshold after multiple hypothesis correction (i.e., *q* ≫ 0.05).

### Analysis and modelling considerations for high time resolution metabolomic data

Although the correlation analysis presented above is based on daily average metabolite intensities (given the disparate, non-paired time scales for the corresponding biometric/nutritional data points), a linear regression model using time of the day as a predictor of metabolite intensity (see **Methods**) established diurnal effects for 268 metabolite features (**Fig. 4a**). Eighty three metabolites had a time of day effect for both subjects, whereas 87 and 98 metabolites only exhibited a time of day effect for Subject 1 or Subject 2, respectively (**Fig. 4a**). Metabolites exhibiting an effect were then separated into three groups based on which time of day showed the greatest effect (calculated here by median z-scores for each time group) (**Fig. 4b** and **Fig. 4c**). For instance, the upper left panel of **Fig. 4c** shows the subset of metabolites varying the most in the morning for Subject 1. For this group of metabolites (amounting to 104/170 metabolites for Subject 1), most are higher in the morning and then decrease over the afternoon and into the evening. For Subject 2, metabolite deviations were more evenly distributed across the day (**Fig. 4b**). Though it is not currently clear what behaviors or underlying biology drive these distinct and time-dependent patterns, it is possible that they originate from differences in dietary habits, age, or physical activity. While Subject 2 did not collect any data on physical activity using a smartphone application, they noted a lack of organized physical activity (e.g., running, weight lifting, bicycling, etc.) during the period of this study.

**Fig. 4.**
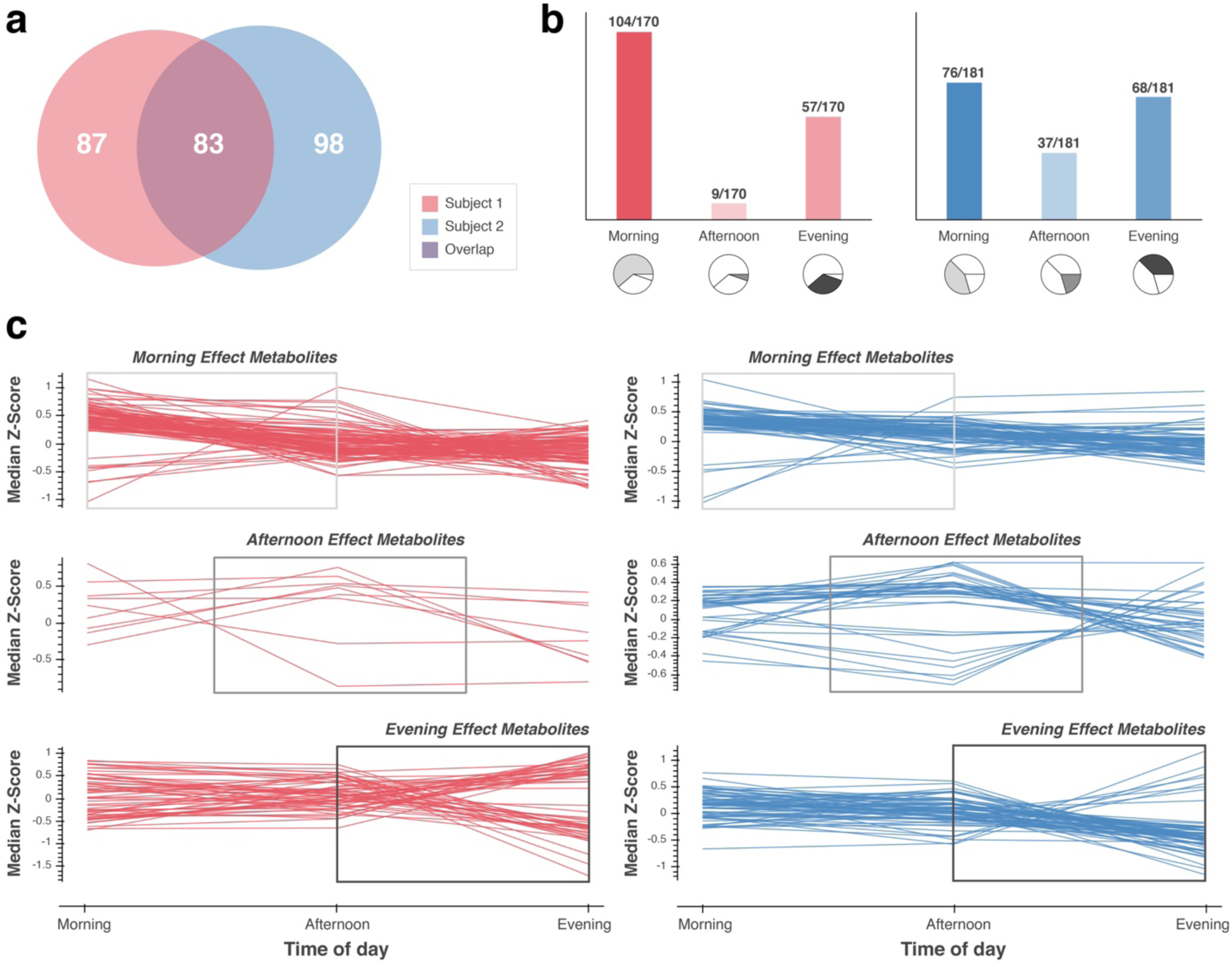
Daily metabolite fluctuations can be shared, distinct, or subject specific. (**a**) A subset of metabolite features for Subject 1 (*n* = 170) and Subject 2 (*n* = 181) were significantly (*p* < 0.05) affected by time of the day based on linear regression analysis (see **Methods**). (**b**) While 83 metabolites were affected by time of the day for both Subjects, 87 and 98 were uniquely affected for Subject 1 and Subject 2, respectively. The majority of metabolites for Subject 1 had the greatest deviation from baseline in the morning, whereas metabolite deviations were more evenly distributed across the day for Subject 2. (**c**) Median Z-scores of metabolite concentrations throughout the course of the day. Most metabolites with the greatest deviations in the morning appear to stabilize by evening (top row), while metabolites with an evening effect (bottom row) are close to baseline concentrations before spiking up or down in the evening. Metabolites with greatest deviations in the afternoon (middle row) peak in the afternoon and tend to reverse course by evening.

Because metabolite concentrations in urine are less directly subjected to the strong homeostatic forces of other biological matrices (such as blood), variance in metabolite concentrations due to diet, lifestyle, and time of the day may be more pronounced^14^. Thus, in light of our analysis, as well as that of other groups^20,38^, there may be better times of the day to measure the effects of factors such as sleep, exercise, or nutrition. Furthermore, optimal sampling time and frequency will likely vary by individual. Thus, at a minimum, it is worth considering diurnal effect (or temporal-behavioral effects) as a confounding variable in more sophisticated modelling approaches with larger, future datasets.

To systematically account for the effect of time dependency and to avoid averaging (and the resulting loss of information, biological variance, and statistical power), we recommend future studies use smartphone applications that record high resolution timestamps for exercise, heart rate, nutritional data, etc. that will allow for more powerful mixed effect linear models^39^. Using a moving average with a smaller interval (i.e., rather than daily average) of metabolite intensity may be another viable approach. However, setting the optimal interval for such a moving average will likely depend on the type of metabolite and the various biological or environmental factors affecting its turnover.

## Discussion

Over the course of 10 days, we measured continuous physiological phenotypes via urine metabolites. Multivariate analysis showed a clear distinction between samples derived from two healthy subjects, suggesting a distinct baseline metabolic fingerprint for each. We observed urine metabolites with known associations to human disease, daily metabolite fluctuations that were subject specific, and saw connections between lifestyle factors such as exercise, nutrition, sleep, and OTC drug usage. Taken together, our data suggest that urine analysis offers metabolic phenotypes that are both quantitative and highly personalized. While healthcare consumer access to such metabolic phenotypes offers tremendous potential for personalized and preventive medicine, both subjects noted practical challenges in participating in this study. For instance, the burden of storing and transporting urine samples using a cooler full of dry ice may explain why few if any other studies have comparable time resolution (i.e., collecting every sample). Future studies and, certainly, user-friendly consumer products would benefit from a collection system that is integrated directly into a toilet.

Of course, designing and manufacturing a consumer-grade device that could effectively and affordably measure metabolites in urine presents many challenges. Such a device should be robust enough to withstand repeated urine analysis from multiple users, sensitive enough to simultaneously quantify tens, hundreds, or even thousands of metabolites, and affordable enough to reduce instead of adding cost to the already overburdened healthcare system. In addition to the technological and economic challenges of building such a device, there are a wide range of ethical challenges in collecting, storing, sharing, and interpreting personalized metabolic information. Though these challenges will hinder development of such a biosensor, we present this dataset and an accompanying interactive web-based visualization tool to share our optimism. We believe that continuous urine metabolite analysis offers a promising opportunity to integrate with current digital technologies as an orthogonal layer of biomedical data to make modern medicine more predictive, preventive, personalized, and participatory.

## Author Contributions

I.J.M., M.S.W., and J.J.C. conceived and planned the study. S.R.P. prepared and analyzed the urine samples using the instrument described in the methods. K.A.O and B.R.P. prepared and analyzed metabolite standards. I.J.M and K.A.O analyzed the data. I.J.M. wrote the code for data analysis and the companion web tool. I.J.M. drafted the manuscript; I.J.M. and J.J.C edited and prepared the manuscript for submission. All authors had access to the data in this study and have read and approve this manuscript.

## Corresponding author

Correspondence to Joshua Coon.

## Acknowledgements

The authors would like to thank Vanessa Linke for useful suggestions regarding analysis and visualization of metabolomics data as well as feedback on the manuscript. The authors would also like to acknowledge Dasom Hwang for her help with refining figures. This research was supported by funding from R35GM118110.

