## Supporting Information - Figures and Tables for "Real time health monitoring through urine metabolomics"

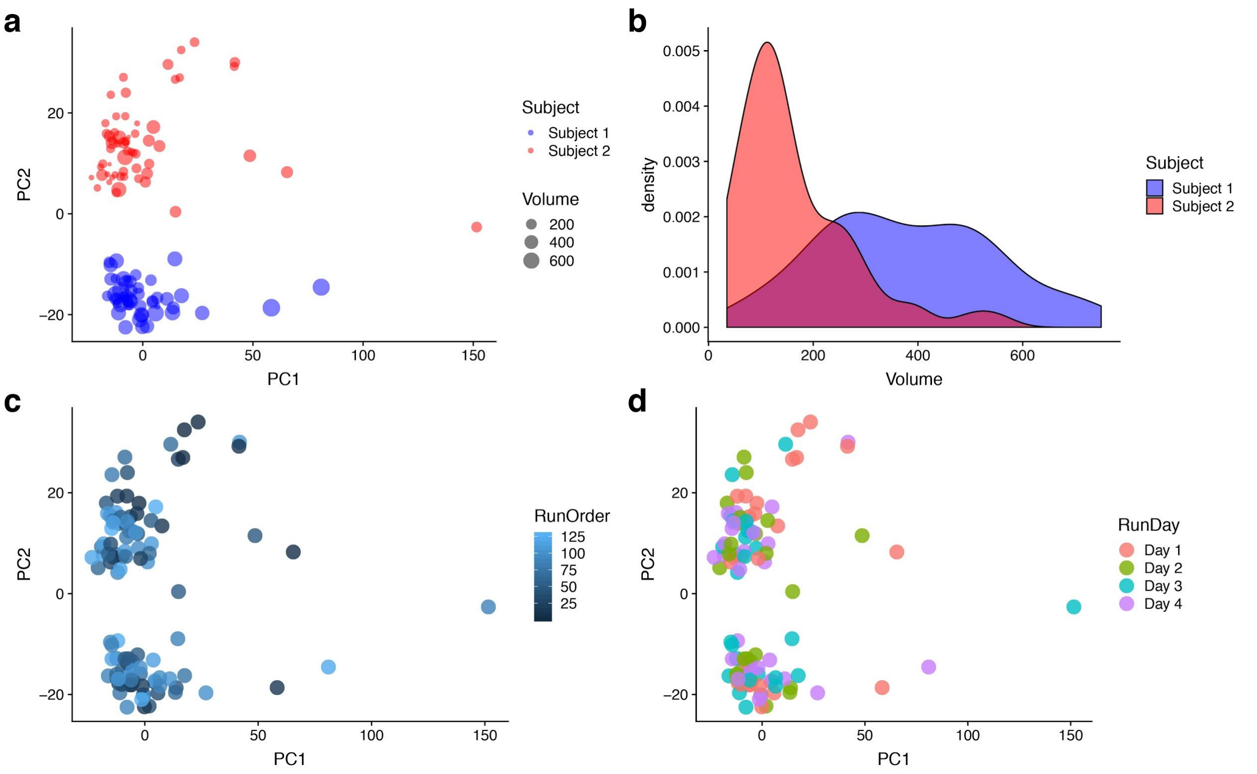


**Supplementary Fig. 1**. PCA-based quality control analysis. (**a**) Samples colored by Subject and sized by volume. (**b**) Density plot of sample volumes for Subject 1 and Subject 2. (**c**) Sample points colored by run order. (**d**) Sample points colored by run day. See **Supplementary Dataset 3** for quantitative values.

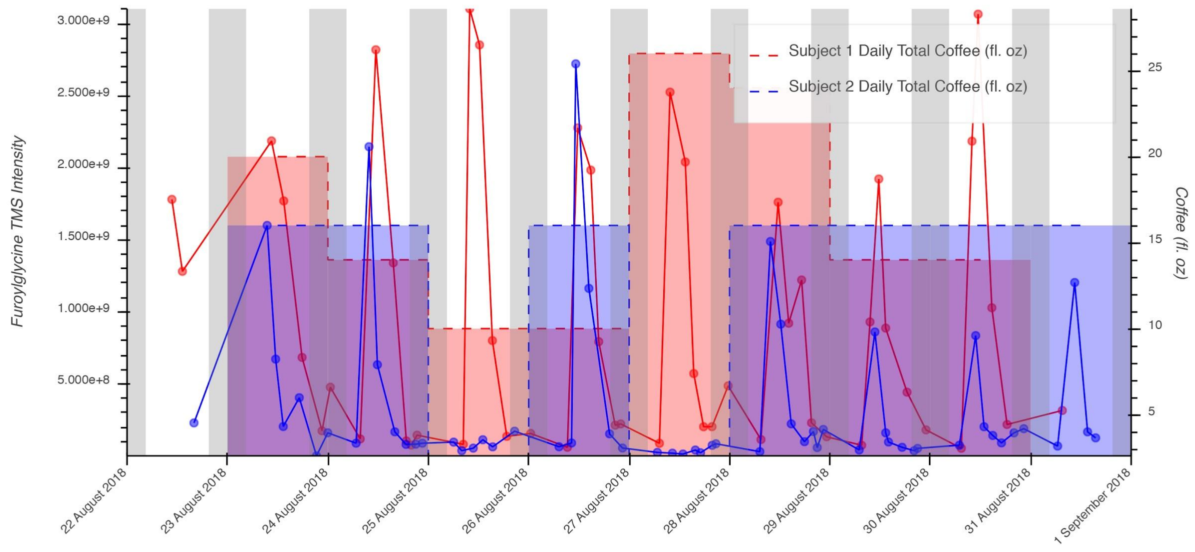


**Supplementary Fig. 2**. Intensity of Furoylglycine TMS over time for both subjects. Coffee (fl. oz) vs. log2-Furoylglycine TMS (daily average intensity); repeated measures r = 0.617, $p$ = 0.011, $q$ = 0.201, dof = 14.

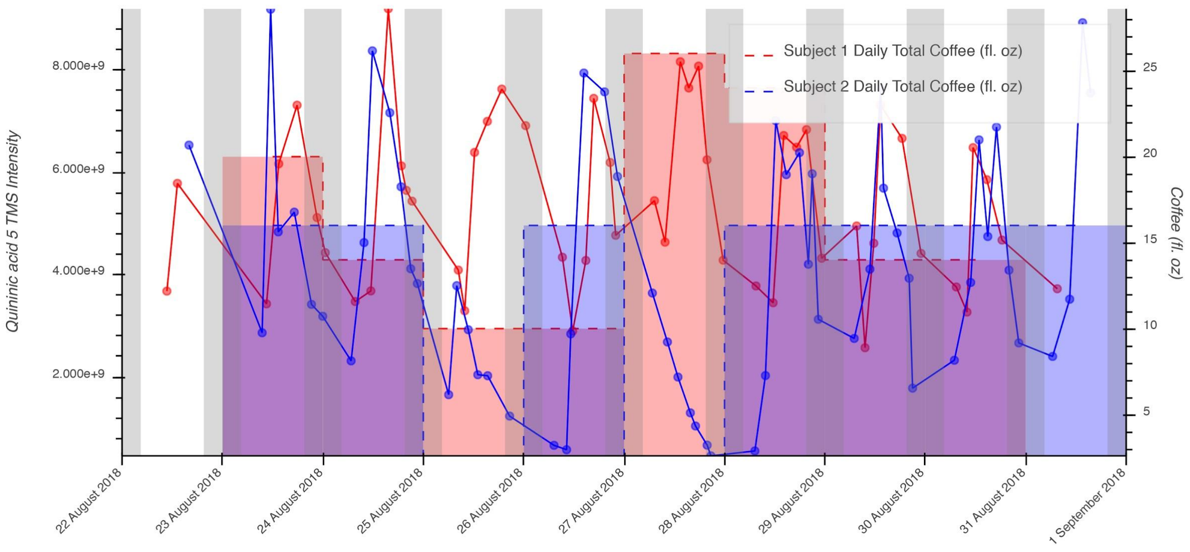


**Supplementary Fig. 3**. Intensity of Quininic acid 5 TMS over time for both subjects. Coffee (fl. oz) vs. log2-Quininic acid 5 TMS (daily average intensity); repeated measures r = 0.787, $p$ = 2.93e-4, $q$ = 0.0884, dof = 14.

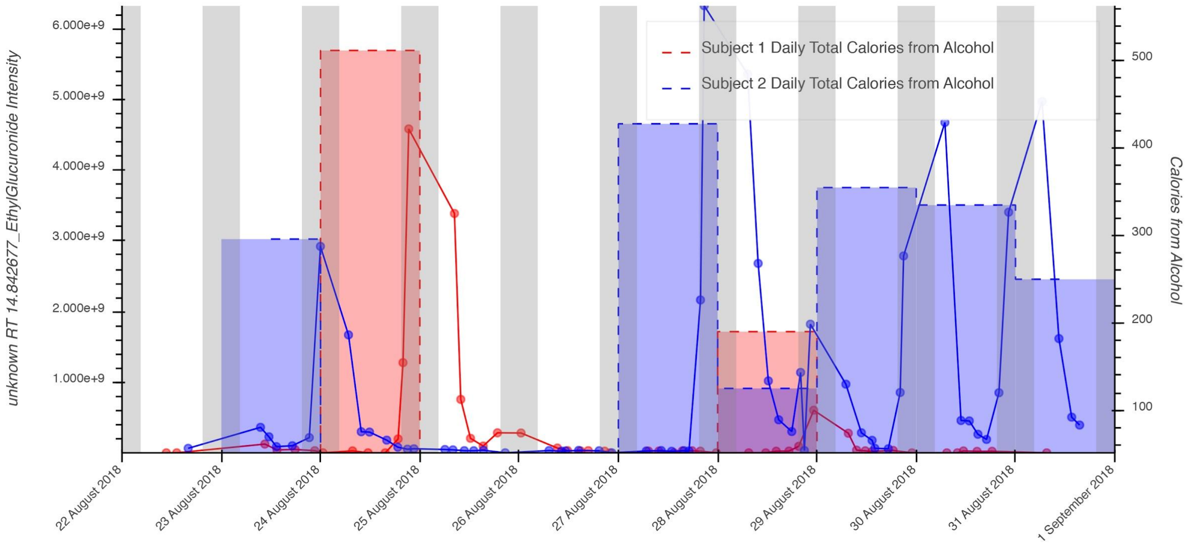


**Supplementary Fig. 4**. Intensity of ethyl glucoronide over time for both subjects. Calories from Alcohol vs. log2-ethyl glucoronide (daily average intensity); repeated measures r = 0.657, $p$ = 0.006, $q$ = 0.0508, dof = 14.

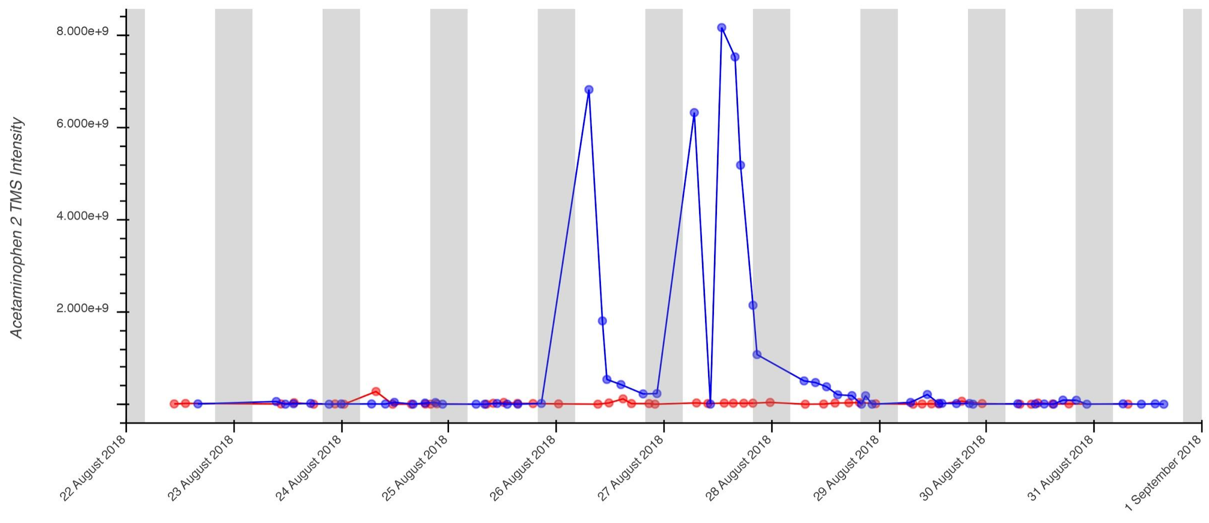


**Supplementary Fig. 5**. Intensity of Acetaminophen 2 TMS over time for both subjects. No quantitative data were recorded for acetaminophen consumption, so correlation metrics are not available.

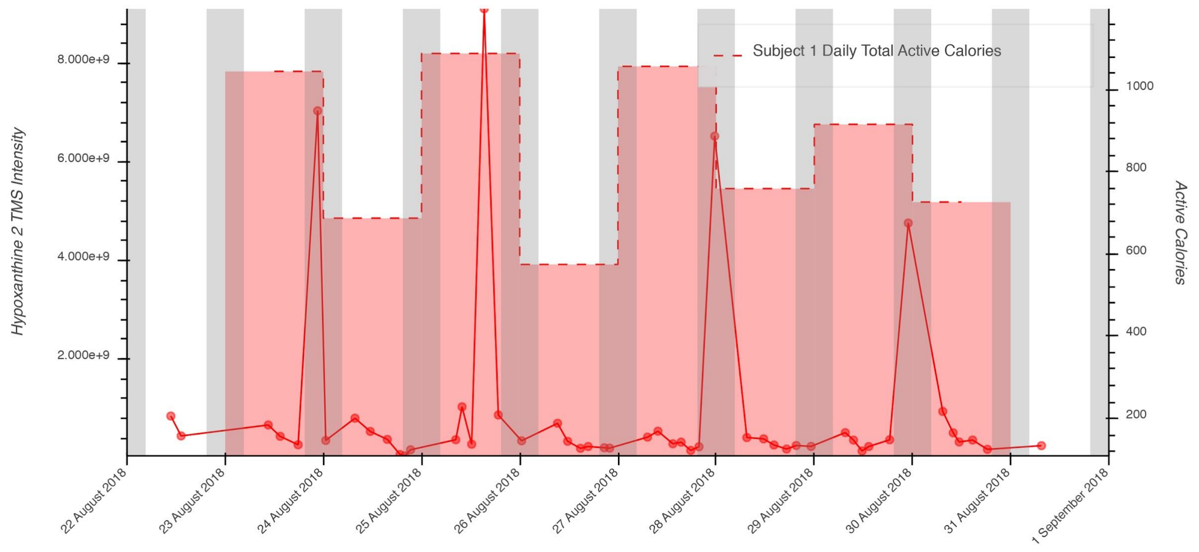


**Supplementary Fig. 6**. Intensity of Hypoxanthine 2 TMS over time for both subjects. Active Calories vs. log2-Hypoxanthine 2 TMS (daily average intensity); Spearman's Rho: 0.833, $p$ = 0.0102, $q$ = 0.472, $n$ = 8.

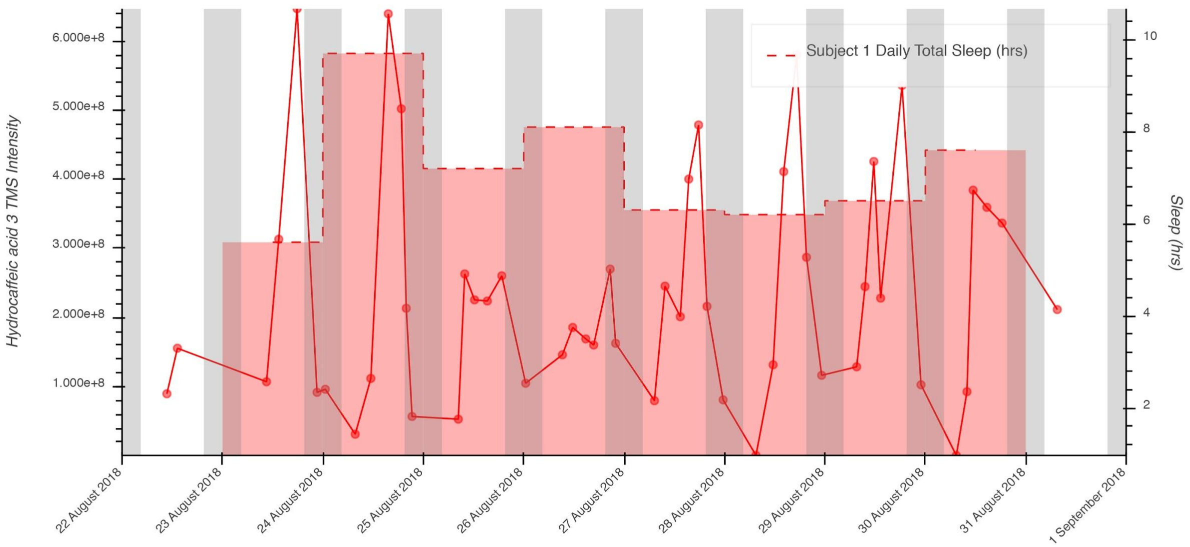


**Supplementary Fig. 7**. Intensity of hydrocaffeic acid 3 TMS over time for both subjects. Sleep (hrs) vs. log2-Hydrocaffeic acid 3 TMS (daily average intensity); Spearman's Rho: -0.857, $p$ = 0.0137, q = 0.551, $n$ = 8.

**Supplementary Table 1**. Summary statistics for biometric measurements.

| **Daily Mean ± (StDev)** | **Subject 1** | **Subject 2** | **App/Hardware** |
|---|---|---|---|
| Dietary Calories (kcal) | 2,904 ± (516) | 1,793 ± (448) | Lose It! |
| Carbohydrates (g) | 295 ± (60) | 146 ± (37) | Lose It! |
| Fiber (g) | 30 ± (15) | 32 ± (11) | Lose It! |
| Sugar (g) | 75 ± (33) | 83 ± (37) | Lose It! |
| Total Fat (g) | 120 ± (29) | 63 ± (25) | Lose It! |
| Saturated Fat (g) | 46 ± (17) | 26 ± (12) | Lose It! |
| Protein (g) | 143 ± (38) | 66 ± (27) | Lose It! |
| Cholesterol (mg) | 352 ± (214) | 180 ± (112) | Lose It! |
| Sodium (mg) | 3,372 ± (983) | 2,063 ± (790) | Lose It! |
| Active Calories (kcal) | 916 ± (214) | NA | Apple Watch |
| Sleep (hrs) | 7.1 (1.2) | NA | Sleep Cycle |

**Supplementary Dataset 1**. Excel file with annotated metabolites and associated diseases from HMDB (hmdb_info.xlsx).

**Supplementary Dataset 2**. Excel file for HMDB disease associations for all available urine metabolites. (hmdb_disease_metabolite_count.xslx).

**Supplementary Dataset 3.** Excel file with combined data for metabolites and samples (combined_sample_data.xlsx).

**Supplementary Dataset 4.** Excel file with correlation analysis results (biometric_correlations.xlsx).
